# Population genetic analysis of 12 X-STRs in a Bahraini population sample (Allele and haplotype frequencies of 12 X-STRs in Bahraini population)

**DOI:** 10.1101/790915

**Authors:** Noora R. Al-Snan, Safia A. Messaoudi, Moiz Bakhiet, Latifa A. Mansoor

**Affiliations:** Department of Molecular Medicine, College of Medical and Medicine Sciences, Arabian Gulf University, Kingdom of Bahrain; Forensic Science Laboratory, Directorate of Forensic Science, General Directorate of Criminal Investigation and Forensic Science, Ministry of Interior, Kingdom of Bahrain; Forensic Sciences Department, College of Criminal Justice, Naif Arab University for Security Sciences, Riyadh, Saudi Arabia

**Keywords:** Bahraini population, Investigator Argus X-12 QS Kit, Forensic parameters, Population genetics, X-STR

## Abstract

To date, there are very limited genetic studies conducted on the Kingdom of Bahrain and this is one of the first studies conducted to evaluate the 12 X-STRs included in the Investigator X-12 QS kit. Bahrain is a small archipelago located in the Arabian Gulf. X-STRs are very informative in population genetics studies, human identification and complex kinship analysis. One hundred and fifty-six (156) buccal swabs were collected using cotton from non-relatives’ Bahraini males from four different regions of Bahrain. DNA was processed using QIAsymphony SP instrument following quantification with Investigator Quantiplex HYres Kit in the 7500 Real-Time PCR System and detected in ABI 3500xl Genetic Analyzer. Analysis was done using different statistical softwares to obtain allele frequencies and haplotype frequencies-based upon the available four clusters of Linkage Groups (LGs)- and the forensic parameters.. Results indicated the diversity of the Bahraini population in terms of high Power of Discrimination (PD) and Probability of Match (PM) values obtained. The combined values of each forensic parameter such as cPD_M_, Mean exclusion chance (CMEC) Krüger, _C_MEC Kishida, and _C_MEC Desmarais as well as _C_MEC Desmarais Duo, were 0.9999983, 9999979, 0.9999939, 0.9999996 and 0.9999514 respectively based on the allele and haplotype frequencies. No shared profiles were observed. Number of non-standard alleles and null samples were obtained with more profoundly in locus DXS10148. We have conducted phylogenetic tree as well as multidimensional scale to analyze the Interpopulation diversity between Bahraini population and nine other populations and found out they reflect the geographical and social background of the region. Overall, the results confirm the importance of X-STRs in discriminating between individuals among Bahraini population and in establishing DNA databases for forensic and kinship studies.

## Introduction

The Investigator Argus X-12 QS Kit (Qiagen, Germany) is a multiplex solution that amplifies 12 segments of DNA located in the X-chromosome; chromosomal X short tandem repeats (X-STRs). This solution is used for human identification, paternity testing, forensic cases and population genetic studies **(1)**. X-chromosome STR markers are very informative for some cases where autosomal STRs fail to provide crucial answers such as in in kinship testing involving female offspring sharing the same father **(2)**. In this kit, the 12 X-STR markers are clustered into four distinct linkage groups (LG), each cluster includes three markers: LG1 (DX8378-DXS10135-DXS10148), LG2 (DXS7132-DXS100740-DXS10079), LG3 (DXS10101-DXS10103-HPRTB), LG4 (DXS7423-DXS10134-DXS10146) **(3)**. The dependency between markers is known as linkage and is the physical proximity of two loci on the same chromosome **(2)**.

To date, there are very limited genetic studies conducted on the Kingdom of Bahrain and knowledge of any such structure is important in the interpretation of the significance of DNA-based forensic evidence and in the construction of appropriate databases. Recent studies was done using autosomal STR and YSTR kits **(4, 5)**.

Kingdom of Bahrain is a small archipelago in total landmass of 760 square kilometers **(6)** consisting of 33 islands, only the five largest are inhabited. These islands are Bahrain, Muharraq, Umm and Nasan and Sitra. Bahrain is positioned in the Arabian Gulf. To the southeast of Bahrain is the State of Qatar, and to its west lies the Kingdom of Saudi Arabia, with which it is connected by a 25-kilometer causeway. To the north and east of Bahrain lies the Islamic Republic of Iran **(7)**.

Because of the geographic location of Bahrain, the diversity of the population had been affected. The Bahraini population is mainly divided into four main ethnic groups: Arabs, Baharna and Persians (Huwala and Ajam) **(8-10)**. This geographical and social organization might be expected to have an expect on the patterns of a genetic diversity **(11)**. Currently, the geographical distribution of Bahraini is included into 4 governorates: Northern, Muharraq, Capital and Southern Governorates.

This present study is to genetically characterize the Bahraini population, using Investigator Argus X-12 QS Kit (Qiagen, Germany). Twelve X-STRs were studied to characterize different genetic population and forensic parameters in 156 Bahraini males. These includes the following primers to co-amplify the Amelogenin (AM) used for gender determination, DXS7132, DXS7423, DXS8378, DXS10074, DXS10079, DXS10101, DXS10103, DXS10134, DXS10135, DXS10146, DXS10148 and HPRTB which have been proven to provide reliable DNA typing results and enhance the power of discrimination (PD). Furthermore, the autosomal STR marker D21S11 is included as a consistent marker for common database kits and prevents sample mixup and excluded of all the genetic analysis for this study. The Investigator Argus X-12 QS Kit Primer Mix contains an internal PCR control (Quality Sensor QS1) to provide helpful information about the efficiency of the PCR. The Quality Sensor is amplified simultaneously with the polymorphic STR markers and was excluded of all the genetic analysis for this study.

## Materials and methods

### Sample Collection

One hundred and fifty-six (156) buccal swabs were collected using cotton swabs (SceneSafe, UK) from non-relatives’ Bahraini males. The research study was announced publicly through different social media channels. Participants who wished to participate contacted the corresponding author for establishing meetings and arrived at the General Directorate of Criminal investigation and Forensic Science – Kingdom of Bahrain to submit their buccal swabs for the research after obtaining informed consent. The age of the participants ranged from 19 to 58 years old.

In each case, males with ancestry (to the level of paternal grandfather) from four different geographical subdivisions of the country (Capital Governorate, Muharraq Governorate, Northern Governorate and Southern Governorate) were sampled. Ethical review for analysis was provided and approved by the Research and Research Ethics Committee (RREC) (E007-PI-10/17) in the Arabian Gulf University. All participants provided informed consent for contribution their buccal swab samples.

### DNA processing

DNAs were extracted using QIAsymphony SP instrument (Qiagen, Germany) following magnetic beads principal. Subsequently the extracted DNAs were quantified using Investigator Quantiplex HYres Kit (Qiagen, Germany) in the 7500 Real-Time PCR System (Thermo Fisher Scientific, Inc., Waltham, MA, USA) according to manufacturer’s recommendation. 0.5 ng of the extracted DNA was amplified using Investigator Argus-X12 QS kit (Qiagen, Germany) with half-volume reactions (12.5µl) following manufactures protocol in 27 cycles. Samples were amplified using MicroAmp Optical 96-Well Reaction Plate (Thermo Fisher Scientific, Inc., Waltham, MA, USA) along with the provided positive control and DNA grade water as a negative control.

The PCR products (1µl) were separated by capillary electrophoresis in an ABI 3500xl Genetic Analyzer (Thermo Fisher Scientific Company, Carlsbad, USA) with reference to the BTO size standard (Qiagen, Germany) in total of 12 µl of BTO size standard and Hi-Di formamide (Thermo Fisher Scientific, Inc., Waltham, MA, USA) master mix. GeneMapper® ID-X Software v1.4 (Thermo Fisher Scientific, Inc., Waltham, MA, USA) was used for genotype assignment.

### Statistical analysis

Allele frequencies and haplotype frequencies were calculated using FreX V1.1 software **(12)**. Also, Linkage groups (LG) were designated as 3 loci in each LG in total of 4 LGs in FreX V1.1 **(12)**. Forensic parameters such as power of discrimination (PD), random matching probability (PM), power of exclusion (PE), polymorphism information content (PIC), number of alleles (Nall) and linkage disequilibrium (LD) between pair of loci were estimated using STRAF - a convenient online tool for STR data evaluation in forensic genetics **(13)**. Mean exclusion chance in Duos (MECD), PD for females (PDf), and PD in males (PDm) were estimated using ChrX-STR.org 2.0 website **(14)**. It should be noted that all of the samples were compromised of males, so it was impossible to calculate the Hardy-Weinberg (HW) equilibrium.

Interpopulation pairwise genetic distances based on Fst between central region Saudi population and the rest of populations extracted from the literature were calculated using POPTREE2 software **(15)** and represented by a nonmetric multidimensional scaling (NM-MDS) analysis using IBM SPSS Statistics v21.0 Software to investigate the populations structure between Bahraini population and the abovementioned populations based on FST’s genetic distances. Phylogenetic tree was constructed from allele frequency data by using the neighbour-joining method **(16)** *via* MEGA X: Molecular Evolutionary Genetics Analysis **(17)**.

It is used to compare between different genetic structure of the populations with Bahraini population using the minimum available loci for different populations. The tree was constructed with allele frequency data of twelve STR loci (DXS7132, DXS7423, DXS8378, DXS10074, DXS10079, DXS10101, DXS10103, DXS10134, DXS10135, DXS10146, DXS10148 and HPRTB) for all populations in corrected fixation index (FST) using neighbor joining for phylogeny in 1000 permutations.

## Results

### Allele frequencies, forensic parameters and efficiency

As for the allele frequency score, some alleles show very high frequencies in the Bahrain population such as allele 14 in DXS7423 with highest frequency of 0.462, followed by allele 19 in DXS10103 with frequency of 0.447 **(Table 1)**. The lowest allele frequency was 0.0064 for 12 different alleles. The probability that two randomly chosen person have the same unspecified genotype at a locus is the sum squares of the frequencies of all genotypes at that locus. The full set of Bahrain data is available in **Table S1**. No shared profiles were observed.

**Table 1.**
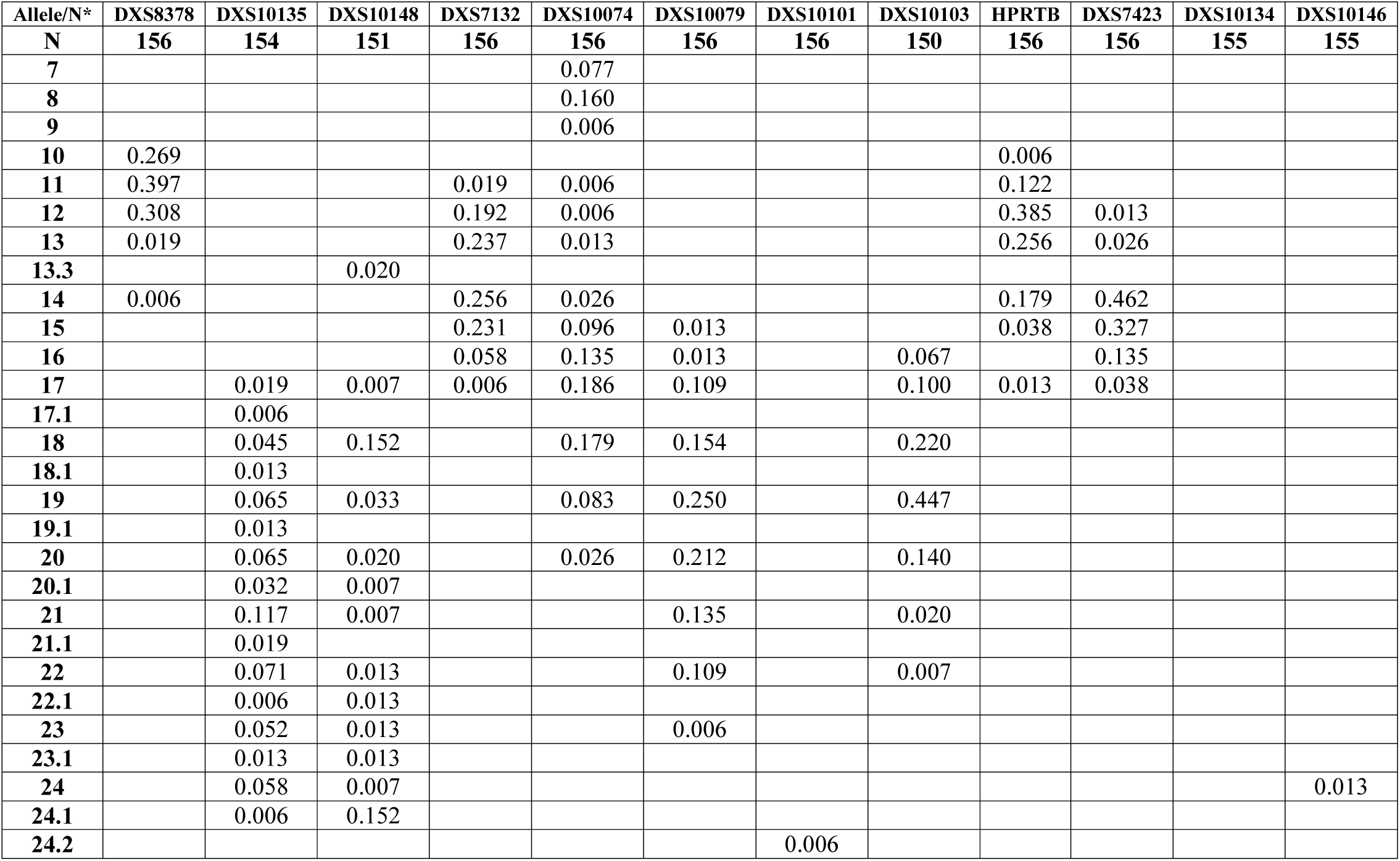
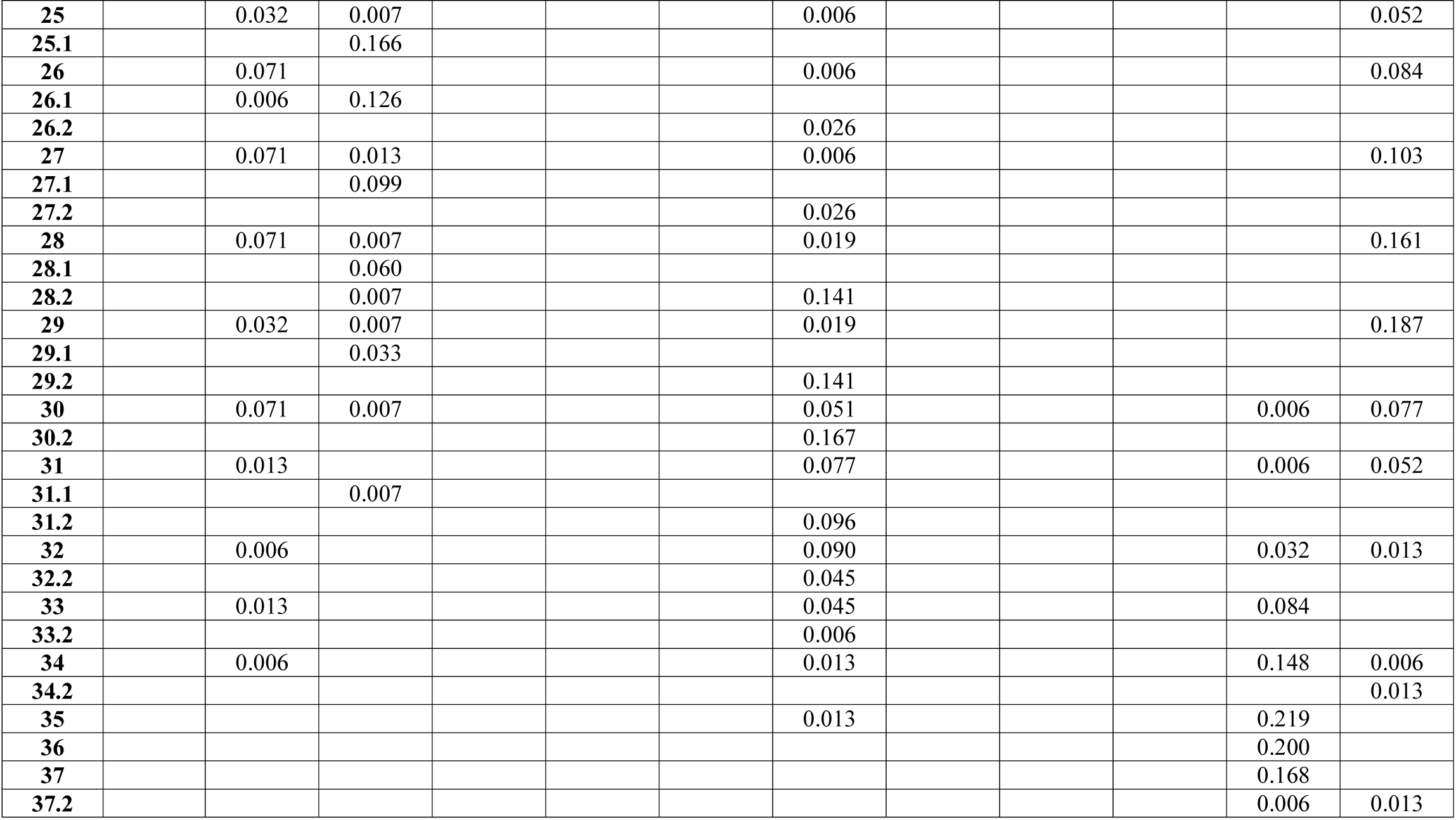
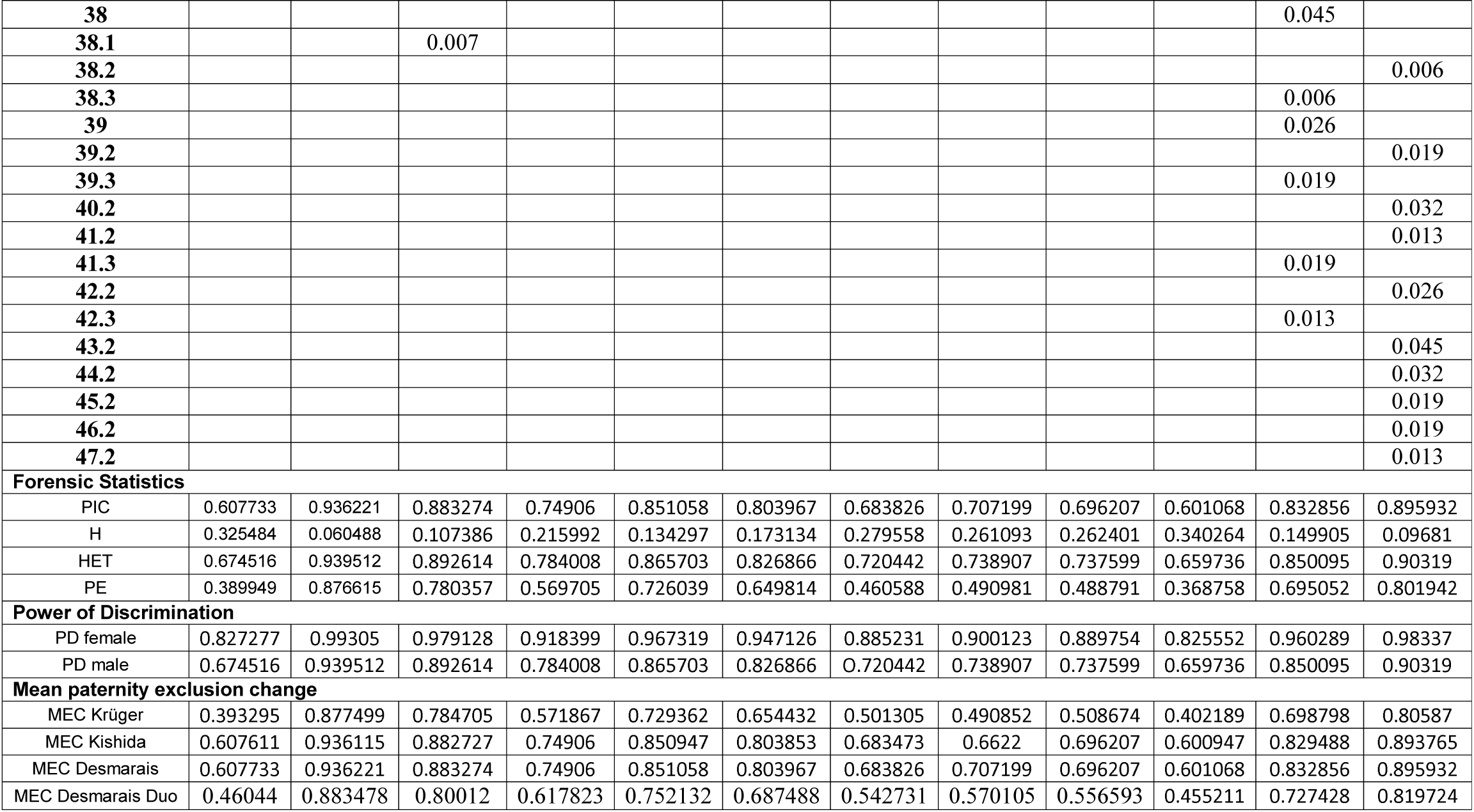
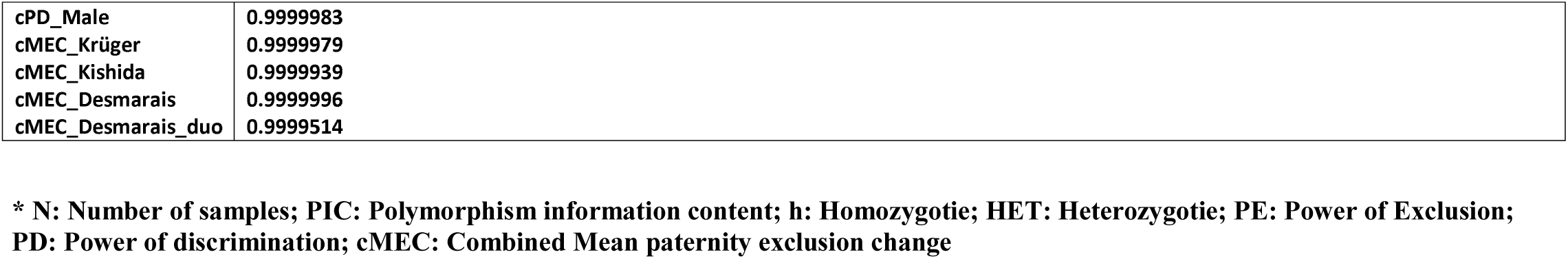
Allele frequency and forensic efficiency parameters of (156) samples from population of Bahrain

| Allele/N* | DXS8378 | DXS10135 | DXS10148 | DXS7132 | DXS10074 | DXS10079 | DXS10101 | DXS10103 | HPRTB | DXS7423 | DXS10134 | DXS10146 |
| --- | --- | --- | --- | --- | --- | --- | --- | --- | --- | --- | --- | --- |
| N | 156 | 154 | 151 | 156 | 156 | 156 | 156 | 150 | 156 | 156 | 155 | 155 |
| 7 |  |  |  |  | 0.077 |  |  |  |  |  |  |  |
| 8 |  |  |  |  | 0.160 |  |  |  |  |  |  |  |
| 9 |  |  |  |  | 0.006 |  |  |  |  |  |  |  |
| 10 | 0.269 |  |  |  |  |  |  |  | 0.006 |  |  |  |
| 11 | 0.397 |  |  | 0.019 | 0.006 |  |  |  | 0.122 |  |  |  |
| 12 | 0.308 |  |  | 0.192 | 0.006 |  |  |  | 0.385 | 0.013 |  |  |
| 13 | 0.019 |  |  | 0.237 | 0.013 |  |  |  | 0.256 | 0.026 |  |  |
| 13.3 |  |  | 0.020 |  |  |  |  |  |  |  |  |  |
| 14 | 0.006 |  |  | 0.256 | 0.026 |  |  |  | 0.179 | 0.462 |  |  |
| 15 |  |  |  | 0.231 | 0.096 | 0.013 |  |  | 0.038 | 0.327 |  |  |
| 16 |  |  |  | 0.058 | 0.135 | 0.013 |  | 0.067 |  | 0.135 |  |  |
| 17 |  | 0.019 | 0.007 | 0.006 | 0.186 | 0.109 |  | 0.100 | 0.013 | 0.038 |  |  |
| 17.1 |  | 0.006 |  |  |  |  |  |  |  |  |  |  |
| 18 |  | 0.045 | 0.152 |  | 0.179 | 0.154 |  | 0.220 |  |  |  |  |
| 18.1 |  | 0.013 |  |  |  |  |  |  |  |  |  |  |
| 19 |  | 0.065 | 0.033 |  | 0.083 | 0.250 |  | 0.447 |  |  |  |  |
| 19.1 |  | 0.013 |  |  |  |  |  |  |  |  |  |  |
| 20 |  | 0.065 | 0.020 |  | 0.026 | 0.212 |  | 0.140 |  |  |  |  |
| 20.1 |  | 0.032 | 0.007 |  |  |  |  |  |  |  |  |  |
| 21 |  | 0.117 | 0.007 |  |  | 0.135 |  | 0.020 |  |  |  |  |
| 21.1 |  | 0.019 |  |  |  |  |  |  |  |  |  |  |
| 22 |  | 0.071 | 0.013 |  |  | 0.109 |  | 0.007 |  |  |  |  |
| 22.1 |  | 0.006 | 0.013 |  |  |  |  |  |  |  |  |  |
| 23 |  | 0.052 | 0.013 |  |  | 0.006 |  |  |  |  |  |  |
| 23.1 |  | 0.013 | 0.013 |  |  |  |  |  |  |  |  |  |
| 24 |  | 0.058 | 0.007 |  |  |  |  |  |  |  |  | 0.013 |
| 24.1 |  | 0.006 | 0.152 |  |  |  |  |  |  |  |  |  |
| 24.2 |  |  |  |  |  |  | 0.006 |  |  |  |  |  |

Table 1 Allele frequency and forensic efficiency parameters of (156) samples from population of Bahrain (Cont.)
| Allele/N | DXS8378 | DXS10135 | DXS10148 | DXS7132 | DXS10074 | DXS10079 | DXS10101 | DXS10103 | HPRTB | DXS7423 | DXS10134 | DXS10146 |
| --- | --- | --- | --- | --- | --- | --- | --- | --- | --- | --- | --- | --- |
| **N** | **156** | **154** | **151** | **156** | **156** | **156** | **156** | **150** | **156** | **156** | **155** | **155** |
| **25** |  | 0.032 | 0.007 |  |  |  | 0.006 |  |  |  |  | 0.052 |
| **25.1** |  |  | 0.166 |  |  |  |  |  |  |  |  |  |
| **26** |  | 0.071 |  |  |  |  | 0.006 |  |  |  |  | 0.084 |
| **26.1** |  | 0.006 | 0.126 |  |  |  |  |  |  |  |  |  |
| **26.2** |  |  |  |  |  |  | 0.026 |  |  |  |  |  |
| **27** |  | 0.071 | 0.013 |  |  |  | 0.006 |  |  |  |  | 0.103 |
| **27.1** |  |  | 0.099 |  |  |  |  |  |  |  |  |  |
| **27.2** |  |  |  |  |  |  | 0.026 |  |  |  |  |  |
| **28** |  | 0.071 | 0.007 |  |  |  | 0.019 |  |  |  |  | 0.161 |
| **28.1** |  |  | 0.060 |  |  |  |  |  |  |  |  |  |
| **28.2** |  |  | 0.007 |  |  |  | 0.141 |  |  |  |  |  |
| **29** |  | 0.032 | 0.007 |  |  |  | 0.019 |  |  |  |  | 0.187 |
| **29.1** |  |  | 0.033 |  |  |  |  |  |  |  |  |  |
| **29.2** |  |  |  |  |  |  | 0.141 |  |  |  |  |  |
| **30** |  | 0.071 | 0.007 |  |  |  | 0.051 |  |  |  | 0.006 | 0.077 |
| **30.2** |  |  |  |  |  |  | 0.167 |  |  |  |  |  |
| **31** |  | 0.013 |  |  |  |  | 0.077 |  |  |  | 0.006 | 0.052 |
| **31.1** |  |  | 0.007 |  |  |  |  |  |  |  |  |  |
| **31.2** |  |  |  |  |  |  | 0.096 |  |  |  |  |  |
| **32** |  | 0.006 |  |  |  |  | 0.090 |  |  |  | 0.032 | 0.013 |
| **32.2** |  |  |  |  |  |  | 0.045 |  |  |  |  |  |
| **33** |  | 0.013 |  |  |  |  | 0.045 |  |  |  | 0.084 |  |
| **33.2** |  |  |  |  |  |  | 0.006 |  |  |  |  |  |
| **34** |  | 0.006 |  |  |  |  | 0.013 |  |  |  | 0.148 | 0.006 |
| **34.2** |  |  |  |  |  |  |  |  |  |  |  | 0.013 |
| **35** |  |  |  |  |  |  | 0.013 |  |  |  | 0.219 |  |
| **36** |  |  |  |  |  |  |  |  |  |  | 0.200 |  |
| **37** |  |  |  |  |  |  |  |  |  |  | 0.168 |  |
| **37.2** |  |  |  |  |  |  |  |  |  |  | 0.006 | 0.013 |

Table 1 Allele frequency and forensic efficiency parameters of (156) samples from population of Bahrain (Cont.)
| Allele/N | DXS8378 | DXS10135 | DXS10148 | DXS7132 | DXS10074 | DXS10079 | DXS10101 | DXS10103 | HPRTB | DXS7423 | DXS10134 | DXS10146 |
| --- | --- | --- | --- | --- | --- | --- | --- | --- | --- | --- | --- | --- |
| N | 156 | 154 | 151 | 156 | 156 | 156 | 156 | 150 | 156 | 156 | 155 | 155 |
| 38 |  |  |  |  |  |  |  |  |  |  | 0.045 |  |
| 38.1 |  |  | 0.007 |  |  |  |  |  |  |  |  |  |
| 38.2 |  |  |  |  |  |  |  |  |  |  |  | 0.006 |
| 38.3 |  |  |  |  |  |  |  |  |  |  | 0.006 |  |
| 39 |  |  |  |  |  |  |  |  |  |  | 0.026 |  |
| 39.2 |  |  |  |  |  |  |  |  |  |  |  | 0.019 |
| 39.3 |  |  |  |  |  |  |  |  |  |  | 0.019 |  |
| 40.2 |  |  |  |  |  |  |  |  |  |  |  | 0.032 |
| 41.2 |  |  |  |  |  |  |  |  |  |  |  | 0.013 |
| 41.3 |  |  |  |  |  |  |  |  |  |  | 0.019 |  |
| 42.2 |  |  |  |  |  |  |  |  |  |  |  | 0.026 |
| 42.3 |  |  |  |  |  |  |  |  |  |  | 0.013 |  |
| 43.2 |  |  |  |  |  |  |  |  |  |  |  | 0.045 |
| 44.2 |  |  |  |  |  |  |  |  |  |  |  | 0.032 |
| 45.2 |  |  |  |  |  |  |  |  |  |  |  | 0.019 |
| 46.2 |  |  |  |  |  |  |  |  |  |  |  | 0.019 |
| 47.2 |  |  |  |  |  |  |  |  |  |  |  | 0.013 |
| **Forensic Statistics** |  |  |  |  |  |  |  |  |  |  |  |  |
| PIC | 0.607733 | 0.936221 | 0.883274 | 0.74906 | 0.851058 | 0.803967 | 0.683826 | 0.707199 | 0.696207 | 0.601068 | 0.832856 | 0.895932 |
| H | 0.325484 | 0.060488 | 0.107386 | 0.215992 | 0.134297 | 0.173134 | 0.279558 | 0.261093 | 0.262401 | 0.340264 | 0.149905 | 0.09681 |
| HET | 0.674516 | 0.939512 | 0.892614 | 0.784008 | 0.865703 | 0.826866 | 0.720442 | 0.738907 | 0.737599 | 0.659736 | 0.850095 | 0.90319 |
| PE | 0.389949 | 0.876615 | 0.780357 | 0.569705 | 0.726039 | 0.649814 | 0.460588 | 0.490981 | 0.488791 | 0.368758 | 0.695052 | 0.801942 |
| **Power of Discrimination** |  |  |  |  |  |  |  |  |  |  |  |  |
| PD female | 0.827277 | 0.99305 | 0.979128 | 0.918399 | 0.967319 | 0.947126 | 0.885231 | 0.900123 | 0.889754 | 0.825552 | 0.960289 | 0.98337 |
| PD male | 0.674516 | 0.939512 | 0.892614 | 0.784008 | 0.865703 | 0.826866 | 0.720442 | 0.738907 | 0.737599 | 0.659736 | 0.850095 | 0.90319 |
| **Mean paternity exclusion change** |  |  |  |  |  |  |  |  |  |  |  |  |
| MEC Krüger | 0.393295 | 0.877499 | 0.784705 | 0.571867 | 0.729362 | 0.654432 | 0.501305 | 0.490852 | 0.508674 | 0.402189 | 0.698798 | 0.80587 |
| MEC Kishida | 0.607611 | 0.936115 | 0.882727 | 0.74906 | 0.850947 | 0.803853 | 0.683473 | 0.6622 | 0.696207 | 0.600947 | 0.829488 | 0.893765 |
| MEC Desmarais | 0.607733 | 0.936221 | 0.883274 | 0.74906 | 0.851058 | 0.803967 | 0.683826 | 0.707199 | 0.696207 | 0.601068 | 0.832856 | 0.895932 |
| MEC Desmarais Duo | 0.46044 | 0.883478 | 0.80012 | 0.617823 | 0.752132 | 0.687488 | 0.542731 | 0.570105 | 0.556593 | 0.455211 | 0.727428 | 0.819724 |

**Table 1 Allele frequency and forensic efficiency parameters of (156) samples from population of Bahrain (Cont.)**
|  |  |
| --- | --- |
| cPD_Male | 0.9999983 |
| cMEC_Krüger | 0.9999979 |
| cMEC_Kishida | 0.9999939 |
| cMEC_Desmarais | 0.9999996 |
| cMEC_Desmarais_duo | 0.9999514 |

**\* N: Number of samples; PIC: Polymorphism information content; h: Homozygotie; HET: Heterozygotie; PE: Power of Exclusion; PD: Power of discrimination; cMEC: Combined Mean paternity exclusion change**

In this studied population, number of alleles per locus was ranged from 5 for marker DXS8378 to 27 for DXS10135. The mean number of alleles per locus was 13.66, and a total number of alleles observed was 164. The most polymorphic locus was DXS10135 **(Table 1)**.

The highest gene diversity (GD) was observed for locus DXS10135 with 0.9467 while the smallest observed was for locus DXS7423 with 0.6693 **(Table 1)**.

Based on allele frequencies, we further determined the statistical parameters of forensic interest. Generally, the polymorphism degree of a specific locus can be measured by the Polymorphism Information Content (PIC) which indicates the degree of genetic polymorphism. We have found out that PIC values for all STR loci were highly informative (PIC ≥ 0.6) with an average of 78.9% **(Table 1)**. PM was ranged from 0.0594 for DXS10135 to 0.3403 for DXS7423.

The power of discrimination (PD) explains level of discriminating between members. The PM was ranged from 0.0594 for DXS10135 to 0.3403 for DXS7423. The PD explains the level of discriminating between members of the same population. As higher the discriminating power of a locus, the more efficient it can be used to discriminate between members. DXS10135 showed the greatest (PD) in Bahraini population with value of 0.9406, whereas DXS7423 gave the lowest value of (PD) with only 0.6597. The average (PD) of the tested loci was 0.815. The combined PD(CPD) and MPCMP) for all the 12 X-STR loci were 99.99999997% and 3.02583E-10 respectively. The combined values of each forensic parameter, cPDM, CMEC Krüger, CMEC Kishida, and CMEC Desmarais as well as CMEC Desmarais Duo, were 0.9999983, 9999979, 0.9999939, 0.9999996 and 0.9999514 respectively **(Table 1)**. The high values of the above-mentioned parameters indicated the usefulness of using the 12XTRs markers included in the Investigator Argus X-12 QS Kit as a discrimination tool in complement with autosomal STRs (4) for genetic characterization of the Bahraini population to differentiate between individuals for forensic and kinship purposes.

### Linkage disequilibrium analysis

As shown in **Table 2**, the study showed no significant deviation from linkage disequilibrium (LD) between pairwise STR loci after Bonferroni’s correction (p > 0.0032 as 0.05/156) in Bahraini population except when plotting the following loci; DXS10074 - DXS10135, DXS10148 - DXS10148, DXS10134 - DXS10148, DXS7132 - DXS10146, DXS7132 - DXS7132 and DXS10103 – HPRTB. The highest pairwise LD was 0.999 when plotting DXS10101 with DXS10148 and also when plotting DXS10103 with DXS10146 and the lowest pairwise LD was 3.62E-07 when plotting DXS10148 with DXS10148.

**Table 2.** Pairwise linkage disequilibrium exact test for all pairs of loci in Bahraini male individuals (n=156)

|  |  |  |  |  |  |  |  |  |  |  |  |
| --- | --- | --- | --- | --- | --- | --- | --- | --- | --- | --- | --- |
| **DXS8378** | **DXS10135** | **DXS10148** | **DXS7132** | **DXS10074** | **DXS10079** | **DXS10101** | **DXS10103** | **HPRTB** | **DXS7423** | **DXS10134** | **DXS10146** |
| **DXS10146** | 0.434436 | 0.834526 | 0.000114 | 0.77173 | 0.623602 | 0.617766 | 0.999879 | 0.762272 | 0.798729 | 0.778209 | 0.460807 |
| **DXS10134** | 0.856572 | 0.00014 | 0.974159 | 0.890384 | 0.009892 | 0.978075 | 0.91026 | 0.119297 | 0.99456 | 0.198766 |  |
| **DXS7423** | 0.12909 | 0.712395 | 0.856028 | 0.58943 | 0.633612 | 0.788513 | 0.874481 | 0.046661 | 0.929244 |  |  |
| **HPRTB** | 0.031025 | 0.360825 | 0.838351 | 0.660623 | 0.719482 | 0.86349 | 6.88E-06 | 0.115648 |  |  |  |
| **DXS10103** | 0.600294 | 0.173926 | 0.236435 | 0.762466 | 0.659187 | 0.750817 | 0.005649 |  |  |  |  |
| **DXS10101** | 0.940213 | 0.999954 | 0.803355 | 0.811644 | 0.870909 | 0.472515 |  |  |  |  |  |
| **DXS10079** | 0.245888 | 0.014217 | 0.982419 | 0.491172 | 0.879475 |  |  |  |  |  |  |
| **DXS10074** | 0.001805 | 0.93321 | 0.963231 | 0.254482 |  |  |  |  |  |  |  |
| **DXS7132** | 0.718462 | 0.839586 | 0.000477 |  |  |  |  |  |  |  |  |
| **DXS10148** | 0.517361 | 2.94E-06 |  |  |  |  |  |  |  |  |  |
| **DXS10135** | 0.498013 |  |  |  |  |  |  |  |  |  |  |
| **DXS8378** |  |  |  |  |  |  |  |  |  |  |  |

The significance in LD was obtained between different loci in different LGs and within same LGs.

### Haplotype allele frequencies

Haplotype allele frequencies were determined using the four LG clusters as shown in **Table 3**. The numbers of observed haplotypes in each of the 4 linkage groups-LG1, LG2, LG3, LG4-were 148, 156, 149 and 153, respectively, while the Haplotype Diversity (HD) values were equal to 1.0000. The three most common haplotype for LG1 was 11-21-25.1 displaying 1.99% of haplotype frequency, in LG2 three sets of haplotypes 14-17-19, 14-7-20 and 15-8-19 were observed each with a frequency of 2.56 % LG3 presented haplotypes 29.2-19-12 4.46% and LG4 14-33-29 and 15-37-27 with haplotype frequency 2.63%.

**Table 3.** Haplotype diversities of four X-Chromosomal linkage groups in Bahraini male individuals

| **LG1**<br>**Haplotype** | **Count** | **Frequency** | **LG2**<br>**Haplotype** | **Count** | **Frequency** | **LG3**<br>**Haplotype** | **Count** | **Frequency** | **LG4**<br>**Haplotype** | **Count** | **Frequency** |
| --- | --- | --- | --- | --- | --- | --- | --- | --- | --- | --- | --- |
| 10 17 18 | 1 | 0.0068 | 11 15 21 | 1 | 0.0064 | 24.2 20 12 | 1 | 0.0067 | 12 37 26 | 1 | 0.0065 |
| 10 17.1 13.3 | 1 | 0.0068 | 11 16 17 | 1 | 0.0064 | 25 19 13 | 1 | 0.0067 | 12 39.3 28 | 1 | 0.0065 |
| 10 19 25.1 | 1 | 0.0068 | 11 16 22 | 1 | 0.0064 | 26 20 15 | 1 | 0.0067 | 13 33 42.2 | 1 | 0.0065 |
| 10 19 26.1 | 1 | 0.0068 | 12 15 18 | 1 | 0.0064 | 26.2 16 12 | 1 | 0.0067 | 13 34 29 | 1 | 0.0065 |
| 10 19.1 26.1 | 1 | 0.0068 | 12 15 19 | 1 | 0.0064 | 26.2 18 12 | 1 | 0.0067 | 13 37 28 | 1 | 0.0065 |
| 10 20 24.1 | 1 | 0.0068 | 12 15 20 | 1 | 0.0064 | 26.2 18 13 | 1 | 0.0067 | 13 39 30 | 1 | 0.0065 |
| 10 20 25.1 | 1 | 0.0068 | 12 15 22 | 1 | 0.0064 | 26.2 20 12 | 1 | 0.0067 | 14 31 32 | 1 | 0.0065 |
| 10 20 26.1 | 1 | 0.0068 | 12 16 19 | 2 | 0.0128 | 27 19 13 | 1 | 0.0067 | 14 32 28 | 2 | 0.0131 |
| 10 20 27.1 | 1 | 0.0068 | 12 16 20 | 1 | 0.0064 | 27.2 18 12 | 1 | 0.0067 | 14 32 29 | 1 | 0.0065 |
| 10 20.1 27.1 | 2 | 0.0135 | 12 16 21 | 1 | 0.0064 | 27.2 21 12 | 1 | 0.0067 | 14 32 31 | 1 | 0.0065 |
| 10 21 18 | 1 | 0.0068 | 12 17 17 | 2 | 0.0128 | 27.2 22 12 | 1 | 0.0067 | 14 33 26 | 1 | 0.0065 |
| 10 21 24.1 | 2 | 0.0135 | 12 17 18 | 1 | 0.0064 | 28 18 12 | 1 | 0.0067 | 14 33 29 | 4 | 0.0261 |
| 10 21 26.1 | 1 | 0.0068 | 12 17 19 | 2 | 0.0128 | 28 19 12 | 1 | 0.0067 | 14 34 25 | 1 | 0.0065 |
| 10 22 27.1 | 1 | 0.0068 | 12 17 21 | 1 | 0.0064 | 28 20 14 | 1 | 0.0067 | 14 34 26 | 3 | 0.0196 |
| 10 23 25.1 | 1 | 0.0068 | 12 18 19 | 1 | 0.0064 | 28.2 16 10 | 1 | 0.0067 | 14 34 28 | 1 | 0.0065 |
| 10 24 26.1 | 1 | 0.0068 | 12 18 20 | 2 | 0.0128 | 28.2 16 12 | 1 | 0.0067 | 14 34 29 | 1 | 0.0065 |
| 10 24 28.1 | 1 | 0.0068 | 12 18 21 | 2 | 0.0128 | 28.2 18 11 | 1 | 0.0067 | 14 34 30 | 2 | 0.0131 |
| 10 24 29.1 | 1 | 0.0068 | 12 19 18 | 1 | 0.0064 | 28.2 18 12 | 1 | 0.0067 | 14 34 31 | 1 | 0.0065 |
| 10 24.1 29.1 | 1 | 0.0068 | 12 19 22 | 1 | 0.0064 | 28.2 18 13 | 3 | 0.0201 | 14 34 41.2 | 2 | 0.0131 |
| 10 25 24.1 | 1 | 0.0068 | 12 7 17 | 1 | 0.0064 | 28.2 19 11 | 5 | 0.0336 | 14 34 44.2 | 1 | 0.0065 |
| 10 25 27.1 | 1 | 0.0068 | 12 7 18 | 1 | 0.0064 | 28.2 19 12 | 3 | 0.0201 | 14 35 24 | 1 | 0.0065 |
| 10 26 19 | 2 | 0.0135 | 12 7 19 | 1 | 0.0064 | 28.2 19 13 | 4 | 0.0268 | 14 35 25 | 3 | 0.0196 |
| 10 26 28.1 | 1 | 0.0068 | 12 7 20 | 1 | 0.0064 | 28.2 20 12 | 1 | 0.0067 | 14 35 26 | 1 | 0.0065 |
| 10 27 17 | 1 | 0.0068 | 12 8 18 | 2 | 0.0128 | 28.2 20 13 | 2 | 0.0134 | 14 35 27 | 1 | 0.0065 |
| 10 27 24.1 | 1 | 0.0068 | 12 8 19 | 1 | 0.0064 | 29 16 14 | 1 | 0.0067 | 14 35 28 | 3 | 0.0196 |
| 10 27 25.1 | 1 | 0.0068 | 12 8 20 | 1 | 0.0064 | 29 19 12 | 1 | 0.0067 | 14 35 29 | 1 | 0.0065 |
| 10 27 26.1 | 1 | 0.0068 | 12 9 18 | 1 | 0.0064 | 29.2 16 13 | 1 | 0.0067 | 14 35 30 | 3 | 0.0196 |
| 10 27 31.1 | 1 | 0.0068 | 13 11 22 | 1 | 0.0064 | 29.2 17 12 | 1 | 0.0067 | 14 35 31 | 2 | 0.0131 |
| 10 28 18 | 1 | 0.0068 | 13 12 20 | 1 | 0.0064 | 29.2 18 11 | 1 | 0.0067 | 14 35 34.2 | 1 | 0.0065 |
| 10 28 23 | 1 | 0.0068 | 13 14 19 | 1 | 0.0064 | 29.2 18 12 | 2 | 0.0134 | 14 35 37.2 | 1 | 0.0065 |
| 10 28 28.1 | 1 | 0.0068 | 13 15 17 | 1 | 0.0064 | 29.2 18 13 | 1 | 0.0067 | 14 35 43.2 | 1 | 0.0065 |
| 10 29 24.1 | 1 | 0.0068 | 13 15 18 | 1 | 0.0064 | 29.2 19 12 | 7 | 0.0470 | 14 35 44.2 | 1 | 0.0065 |
| 10 29 25.1 | 2 | 0.0135 | 13 15 19 | 2 | 0.0128 | 29.2 19 13 | 3 | 0.0201 | 14 35 47.2 | 1 | 0.0065 |
| 10 29 27.1 | 1 | 0.0068 | 13 15 20 | 1 | 0.0064 | 29.2 19 14 | 2 | 0.0134 | 14 36 26 | 1 | 0.0065 |

Table 3 Haplotype diversities of four X-Chromosomal linkage groups in Bahraini male individuals (Cont.)
| LG1<br>Haplotype | Count | Frequency | LG2<br>Haplotype | Count | Frequency | LG3<br>Haplotype | Count | Frequency | LG4<br>Haplotype | Count | Frequency |
| --- | --- | --- | --- | --- | --- | --- | --- | --- | --- | --- | --- |
| 10 30 19 | 1 | 0.0068 | 13 15 22 | 1 | 0.0064 | 29.2 19 15 | 1 | 0.0067 | 14 36 27 | 1 | 0.0065 |
| 10 30 19 | 1 | 0.0068 | 13 15 22 | 1 | 0.0064 | 29.2 19 15 | 1 | 0.0067 | 14 36 27 | 1 | 0.0065 |
| 10 30 25.1 | 1 | 0.0068 | 13 16 15 | 1 | 0.0064 | 29.2 20 12 | 1 | 0.0067 | 14 36 28 | 2 | 0.0131 |
| 10 30 26.1 | 1 | 0.0068 | 13 16 18 | 1 | 0.0064 | 30 16 14 | 1 | 0.0067 | 14 36 29 | 2 | 0.0131 |
| 10 31 25.1 | 1 | 0.0068 | 13 16 20 | 1 | 0.0064 | 30 17 12 | 1 | 0.0067 | 14 36 30 | 1 | 0.0065 |
| 11 17 27 | 2 | 0.0135 | 13 16 21 | 1 | 0.0064 | 30 18 12 | 1 | 0.0067 | 14 36 37.2 | 1 | 0.0065 |
| 11 18 13.3 | 1 | 0.0068 | 13 17 19 | 1 | 0.0064 | 30 19 12 | 2 | 0.0134 | 14 36 42.2 | 2 | 0.0131 |
| 11 18 25.1 | 2 | 0.0135 | 13 17 20 | 1 | 0.0064 | 30 19 13 | 1 | 0.0067 | 14 36 43.2 | 1 | 0.0065 |
| 11 18.1 18 | 2 | 0.0135 | 13 17 21 | 3 | 0.0192 | 30 19 14 | 1 | 0.0067 | 14 36 45.2 | 1 | 0.0065 |
| 11 19 18 | 1 | 0.0068 | 13 17 22 | 1 | 0.0064 | 30.2 18 11 | 4 | 0.0268 | 14 37 27 | 1 | 0.0065 |
| 11 19 25.1 | 2 | 0.0135 | 13 17 23 | 1 | 0.0064 | 30.2 18 12 | 1 | 0.0067 | 14 37 28 | 1 | 0.0065 |
| 11 20 18 | 1 | 0.0068 | 13 18 15 | 1 | 0.0064 | 30.2 18 13 | 1 | 0.0067 | 14 37 29 | 3 | 0.0196 |
| 11 20 26.1 | 1 | 0.0068 | 13 18 18 | 1 | 0.0064 | 30.2 18 14 | 1 | 0.0067 | 14 37 30 | 1 | 0.0065 |
| 11 20.1 24.1 | 1 | 0.0068 | 13 18 19 | 3 | 0.0192 | 30.2 18 15 | 2 | 0.0134 | 14 37 42.2 | 1 | 0.0065 |
| 11 21 18 | 2 | 0.0135 | 13 18 20 | 2 | 0.0128 | 30.2 19 11 | 3 | 0.0201 | 14 37 46.2 | 1 | 0.0065 |
| 11 21 19 | 1 | 0.0068 | 13 18 21 | 1 | 0.0064 | 30.2 19 12 | 4 | 0.0268 | 14 37.2 29 | 1 | 0.0065 |
| 11 21 20 | 1 | 0.0068 | 13 18 22 | 1 | 0.0064 | 30.2 19 13 | 3 | 0.0201 | 14 38 28 | 1 | 0.0065 |
| 11 21 24.1 | 2 | 0.0135 | 13 19 17 | 1 | 0.0064 | 30.2 19 15 | 1 | 0.0067 | 14 38 46.2 | 1 | 0.0065 |
| 11 21 25.1 | 3 | 0.0203 | 13 19 21 | 1 | 0.0064 | 30.2 20 11 | 2 | 0.0134 | 14 38.3 26 | 1 | 0.0065 |
| 11 21 26.1 | 1 | 0.0068 | 13 19 22 | 1 | 0.0064 | 30.2 20 12 | 1 | 0.0067 | 14 39 28 | 2 | 0.0131 |
| 11 21 28.2 | 1 | 0.0068 | 13 8 17 | 1 | 0.0064 | 30.2 20 14 | 2 | 0.0134 | 14 39.3 31 | 1 | 0.0065 |
| 11 21 29.1 | 1 | 0.0068 | 13 8 18 | 1 | 0.0064 | 30.2 21 12 | 1 | 0.0067 | 14 41.3 27 | 1 | 0.0065 |
| 11 21.1 27.1 | 1 | 0.0068 | 13 8 20 | 1 | 0.0064 | 31 16 13 | 1 | 0.0067 | 14 42.3 27 | 1 | 0.0065 |
| 11 22 18 | 1 | 0.0068 | 13 8 21 | 1 | 0.0064 | 31 17 12 | 3 | 0.0201 | 15 30 30 | 1 | 0.0065 |
| 11 22 22 | 1 | 0.0068 | 13 8 22 | 1 | 0.0064 | 31 17 13 | 1 | 0.0067 | 15 33 28 | 1 | 0.0065 |
| 11 22 24.1 | 2 | 0.0135 | 14 13 22 | 1 | 0.0064 | 31 18 12 | 2 | 0.0134 | 15 33 29 | 2 | 0.0131 |
| 11 22 26.1 | 1 | 0.0068 | 14 15 20 | 1 | 0.0064 | 31 18 13 | 1 | 0.0067 | 15 33 31 | 1 | 0.0065 |
| 11 22 30 | 1 | 0.0068 | 14 16 16 | 1 | 0.0064 | 31 19 13 | 1 | 0.0067 | 15 33 43.2 | 1 | 0.0065 |
| 11 22.1 19 | 1 | 0.0068 | 14 16 18 | 1 | 0.0064 | 31 19 14 | 1 | 0.0067 | 15 34 28 | 2 | 0.0131 |
| 11 23 23 | 1 | 0.0068 | 14 16 19 | 3 | 0.0192 | 31 20 12 | 1 | 0.0067 | 15 34 29 | 1 | 0.0065 |
| 11 23 24.1 | 1 | 0.0068 | 14 16 20 | 1 | 0.0064 | 31 20 13 | 1 | 0.0067 | 15 34 40.2 | 1 | 0.0065 |
| 11 23 25.1 | 1 | 0.0068 | 14 16 22 | 2 | 0.0128 | 31.2 16 12 | 1 | 0.0067 | 15 35 25 | 2 | 0.0131 |
| 11 23 26.1 | 1 | 0.0068 | 14 17 18 | 1 | 0.0064 | 31.2 17 14 | 1 | 0.0067 | 15 35 26 | 1 | 0.0065 |

Table 3 Haplotype diversities of four X-Chromosomal linkage groups in Bahraini male individuals (Cont.)
| LG1 Haplotype | Count | Frequency | LG2 Haplotype | Count | Frequency | LG3 Haplotype | Count | Frequency | LG4 Haplotype | Count | Frequency |
| --- | --- | --- | --- | --- | --- | --- | --- | --- | --- | --- | --- |
| 11 23 28 | 1 | 0.0068 | 14 17 19 | 4 | 0.0256 | 31.2 18 12 | 1 | 0.0067 | 15 35 28 | 1 | 0.0065 |
| 11 23.1 25.1 | 1 | 0.0068 | 14 17 21 | 1 | 0.0064 | 31.2 19 12 | 3 | 0.0201 | 15 35 31 | 1 | 0.0065 |
| 11 24 18 | 1 | 0.0068 | 14 18 17 | 1 | 0.0064 | 31.2 19 13 | 4 | 0.0268 | 15 35 34 | 1 | 0.0065 |
| 11 24 20 | 1 | 0.0068 | 14 18 18 | 2 | 0.0128 | 31.2 19 14 | 3 | 0.0201 | 15 35 39.2 | 1 | 0.0065 |
| 11 24 23.1 | 1 | 0.0068 | 14 18 19 | 2 | 0.0128 | 31.2 20 14 | 1 | 0.0067 | 15 35 43.2 | 1 | 0.0065 |
| 11 24 24.1 | 1 | 0.0068 | 14 18 20 | 2 | 0.0128 | 31.2 21 13 | 1 | 0.0067 | 15 35 44.2 | 1 | 0.0065 |
| 11 24 25.1 | 1 | 0.0068 | 14 18 22 | 1 | 0.0064 | 32 16 14 | 1 | 0.0067 | 15 36 24 | 1 | 0.0065 |
| 11 25 27.1 | 1 | 0.0068 | 14 19 18 | 2 | 0.0128 | 32 17 12 | 3 | 0.0201 | 15 36 26 | 1 | 0.0065 |
| 11 26 24.1 | 1 | 0.0068 | 14 19 22 | 1 | 0.0064 | 32 17 14 | 3 | 0.0201 | 15 36 28 | 3 | 0.0196 |
| 11 26 25.1 | 1 | 0.0068 | 14 20 20 | 1 | 0.0064 | 32 18 13 | 2 | 0.0134 | 15 36 29 | 2 | 0.0131 |
| 11 26.1 24 | 1 | 0.0068 | 14 20 22 | 1 | 0.0064 | 32 19 14 | 3 | 0.0201 | 15 36 30 | 1 | 0.0065 |
| 11 27 18 | 1 | 0.0068 | 14 7 17 | 1 | 0.0064 | 32 20 14 | 2 | 0.0134 | 15 36 40.2 | 3 | 0.0196 |
| 11 27 25.1 | 1 | 0.0068 | 14 7 20 | 4 | 0.0256 | 32.2 18 13 | 1 | 0.0067 | 15 36 46.2 | 1 | 0.0065 |
| 11 28 24.1 | 1 | 0.0068 | 14 8 18 | 1 | 0.0064 | 32.2 19 12 | 1 | 0.0067 | 15 37 26 | 2 | 0.0131 |
| 11 28 26.1 | 1 | 0.0068 | 14 8 19 | 3 | 0.0192 | 32.2 19 14 | 2 | 0.0134 | 15 37 27 | 4 | 0.0261 |
| 11 28 29.1 | 1 | 0.0068 | 14 8 20 | 1 | 0.0064 | 32.2 20 12 | 2 | 0.0134 | 15 37 28 | 1 | 0.0065 |
| 11 30 24.1 | 1 | 0.0068 | 14 8 21 | 1 | 0.0064 | 32.2 20 13 | 1 | 0.0067 | 15 37 29 | 2 | 0.0131 |
| 11 30 27.1 | 1 | 0.0068 | 15 13 19 | 1 | 0.0064 | 33 16 13 | 2 | 0.0134 | 15 37 39.2 | 1 | 0.0065 |
| 11 30 28.1 | 1 | 0.0068 | 15 14 17 | 1 | 0.0064 | 33 17 12 | 1 | 0.0067 | 15 37 47.2 | 1 | 0.0065 |
| 11 31 28.1 | 1 | 0.0068 | 15 14 20 | 2 | 0.0128 | 33 17 14 | 1 | 0.0067 | 15 38 27 | 2 | 0.0131 |
| 11 32 22 | 1 | 0.0068 | 15 15 18 | 1 | 0.0064 | 33 19 14 | 1 | 0.0067 | 15 38 29 | 1 | 0.0065 |
| 11 34 26.1 | 1 | 0.0068 | 15 16 18 | 1 | 0.0064 | 33 19 17 | 2 | 0.0134 | 15 38 34.2 | 1 | 0.0065 |
| 12 18 18 | 1 | 0.0068 | 15 16 19 | 1 | 0.0064 | 33.2 19 12 | 1 | 0.0067 | 15 38 39.2 | 1 | 0.0065 |
| 12 18 25 | 1 | 0.0068 | 15 17 16 | 1 | 0.0064 | 34 18 11 | 1 | 0.0067 | 15 39 27 | 1 | 0.0065 |
| 12 18 26.1 | 1 | 0.0068 | 15 17 17 | 1 | 0.0064 | 34 19 13 | 1 | 0.0067 | 15 39.3 30 | 1 | 0.0065 |
| 12 19 18 | 2 | 0.0135 | 15 17 19 | 2 | 0.0128 |  |  |  | 15 42.3 28 | 1 | 0.0065 |
| 12 19 20.1 | 1 | 0.0068 | 15 17 20 | 3 | 0.0192 |  |  |  | 16 33 27 | 1 | 0.0065 |
| 12 19 38.1 | 1 | 0.0068 | 15 17 21 | 2 | 0.0128 |  |  |  | 16 34 29 | 1 | 0.0065 |
| 12 19.1 28.1 | 1 | 0.0068 | 15 17 22 | 1 | 0.0064 |  |  |  | 16 34 30 | 1 | 0.0065 |
| 12 20 18 | 1 | 0.0068 | 15 18 17 | 1 | 0.0064 |  |  |  | 16 34 43.2 | 1 | 0.0065 |
| 12 20 24.1 | 2 | 0.0135 | 15 18 19 | 2 | 0.0128 |  |  |  | 16 35 29 | 2 | 0.0131 |
| 12 20 27.1 | 1 | 0.0068 | 15 18 20 | 1 | 0.0064 |  |  |  | 16 35 32 | 1 | 0.0065 |

**Table 3 Haplotype diversities of four X-Chromosomal linkage groups in Bahraini male individuals (Cont.)**
| **LG1**<br>**Haplotype** | **Count** | **Frequency** | **LG2**<br>**Haplotype** | **Count** | **Frequency** | **LG3**<br>**Haplotype** | **Count** | **Frequency** | **LG4**<br>**Haplotype** | **Count** | **Frequency** |
| --- | --- | --- | --- | --- | --- | --- | --- | --- | --- | --- | --- |
| 12 20.1 18 | 1 | 0.0068 | 15 19 18 | 1 | 0.0064 |  |  |  | 16 35 38.2 | 1 | 0.0065 |
| 12 20.1 22.1 | 1 | 0.0068 | 15 19 19 | 1 | 0.0064 |  |  |  | 16 35 45.2 | 1 | 0.0065 |
| 12 21 26.1 | 1 | 0.0068 | 15 19 21 | 2 | 0.0128 |  |  |  | 16 36 26 | 1 | 0.0065 |
| 12 21 27.1 | 1 | 0.0068 | 15 7 17 | 1 | 0.0064 |  |  |  | 16 36 28 | 2 | 0.0131 |
| 12 21.1 18 | 1 | 0.0068 | 15 7 19 | 1 | 0.0064 |  |  |  | 16 36 29 | 2 | 0.0131 |
| 12 22 13.3 | 1 | 0.0068 | 15 7 22 | 1 | 0.0064 |  |  |  | 16 36 31 | 1 | 0.0065 |
| 12 22 24.1 | 1 | 0.0068 | 15 8 18 | 2 | 0.0128 |  |  |  | 16 37 25 | 1 | 0.0065 |
| 12 23 27.1 | 1 | 0.0068 | 15 8 19 | 4 | 0.0256 |  |  |  | 16 37 27 | 2 | 0.0131 |
| 12 24 27.1 | 1 | 0.0068 | 15 8 21 | 2 | 0.0128 |  |  |  | 16 37 29 | 1 | 0.0065 |
| 12 24 29 | 1 | 0.0068 | 16 15 18 | 1 | 0.0064 |  |  |  | 16 37 44.2 | 1 | 0.0065 |
| 12 25 26.1 | 2 | 0.0135 | 16 16 20 | 1 | 0.0064 |  |  |  | 17 32 44.2 | 1 | 0.0065 |
| 12 26 18 | 2 | 0.0135 | 16 18 17 | 2 | 0.0128 |  |  |  | 17 33 25 | 1 | 0.0065 |
| 12 26 24.1 | 1 | 0.0068 | 16 19 20 | 1 | 0.0064 |  |  |  | 17 34 43.2 | 2 | 0.0131 |
| 12 26 25.1 | 1 | 0.0068 | 16 20 17 | 2 | 0.0128 |  |  |  | 17 34 45.2 | 1 | 0.0065 |
| 12 26 27.1 | 1 | 0.0068 | 16 8 20 | 1 | 0.0064 |  |  |  | 17 41.3 40.2 | 1 | 0.0065 |
| 12 26 28.1 | 1 | 0.0068 | 16 8 21 | 1 | 0.0064 |  |  |  |  |  |  |
| 12 27 18 | 1 | 0.0068 | 17 15 20 | 1 | 0.0064 |  |  |  |  |  |  |
| 12 27 24.1 | 2 | 0.0135 |  |  |  |  |  |  |  |  |  |
| 12 28 25.1 | 2 | 0.0135 |  |  |  |  |  |  |  |  |  |
| 12 28 27.1 | 1 | 0.0068 |  |  |  |  |  |  |  |  |  |
| 12 28 28.1 | 2 | 0.0135 |  |  |  |  |  |  |  |  |  |
| 12 29 25.1 | 1 | 0.0068 |  |  |  |  |  |  |  |  |  |
| 12 29 29.1 | 1 | 0.0068 |  |  |  |  |  |  |  |  |  |
| 12 30 18 | 1 | 0.0068 |  |  |  |  |  |  |  |  |  |
| 12 30 21 | 1 | 0.0068 |  |  |  |  |  |  |  |  |  |
| 12 30 25.1 | 1 | 0.0068 |  |  |  |  |  |  |  |  |  |
| 12 30 26.1 | 1 | 0.0068 |  |  |  |  |  |  |  |  |  |
| 12 33 22.1 | 1 | 0.0068 |  |  |  |  |  |  |  |  |  |
| 12 33 26.1 | 1 | 0.0068 |  |  |  |  |  |  |  |  |  |
| 13 23 18 | 1 | 0.0068 |  |  |  |  |  |  |  |  |  |

LG1 gave the highest number of haplotype combinations (127) whileLG3 gave the lowest number of haplotype combinations (90). As for LG1, most of haplotype frequencies counted as 1 or 2 which gave very diverse combinations of haplotypes. Unique haplotypes, which were observed once, accounted for 55.9% of all LGs haplotype observed (339/606).

### Rare variants, off-ladder and null alleles

Several cases showed off ladder (OL) in various loci; two allelic ladder variants were detected at the DXS10146; Sample#12 indicated OL in 223.05 bp and sample#34 showed OL in 243.87 bp. Four allelic ladder variants were detected at DXS10148; Sample#29 showed OL in 312.62 bp, sample#49 and sample#50 showed OL in 312.57, and sample#123 showed OL in 280.19 bp. One allelic ladder variants were detected with OL in DXS10134; Sample#72 showed OL with 325.61 bp. Null alleles were also observed in two loci; DXS10148 in sample#105 and in DXS10103 showed drop out allele in sample#135 and sample#119 **(Fig 1**).

**Figure 1.**
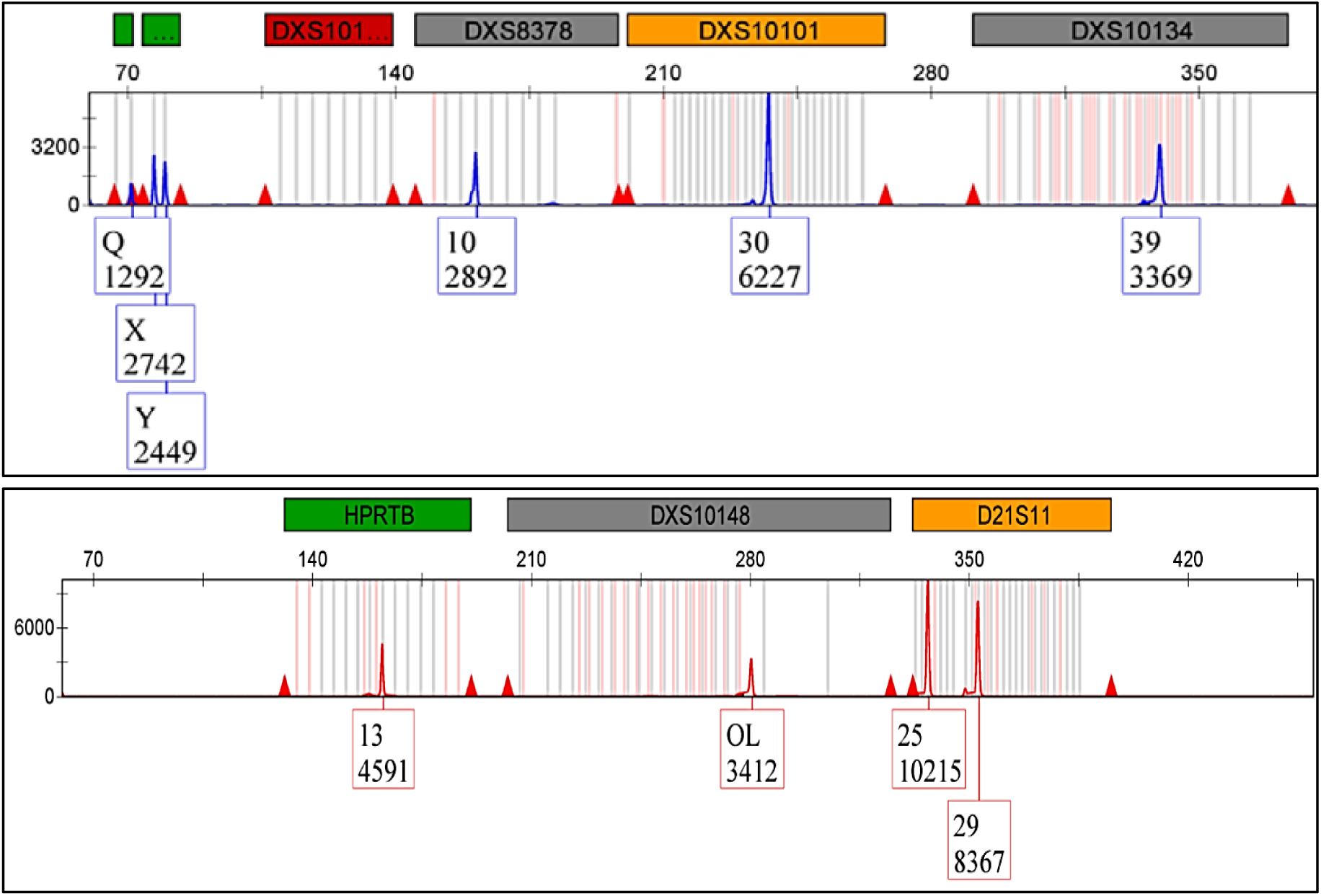
Non standard and null cases observed in some samples

High frequent cases of OL and null alleles were in DXS10148, which gave indication that the Investigator Argus X-12 QS kit provided standard ladder lack significant coverage of bins at DXS10148 as confirmed in previous literature **(18)**.

More than half of the samples (103/156) showed 0.1 variant in DXS10148 and 18 samples gave the same variant in DXS10135.

More than half of the samples (102/156) also showed 0.2 variant in DXS10101, 39 samples in DXS10146, one sample in DXS10134 and one in DXS10148. As for 0.3 variant; 9 samples were detected in DXS10134 and 3 samples in DXS10148.

### Interpopulation diversity

To measure the diversity between Bahraini population and other populations previously reported, we have constructed the phylogenetic tree from allelelic frequencies data by using the neighbor-joining (NJ) method *via* MEGA X: Molecular Evolutionary Genetics Analysis **(Fig 2)**.

**Figure 2.**
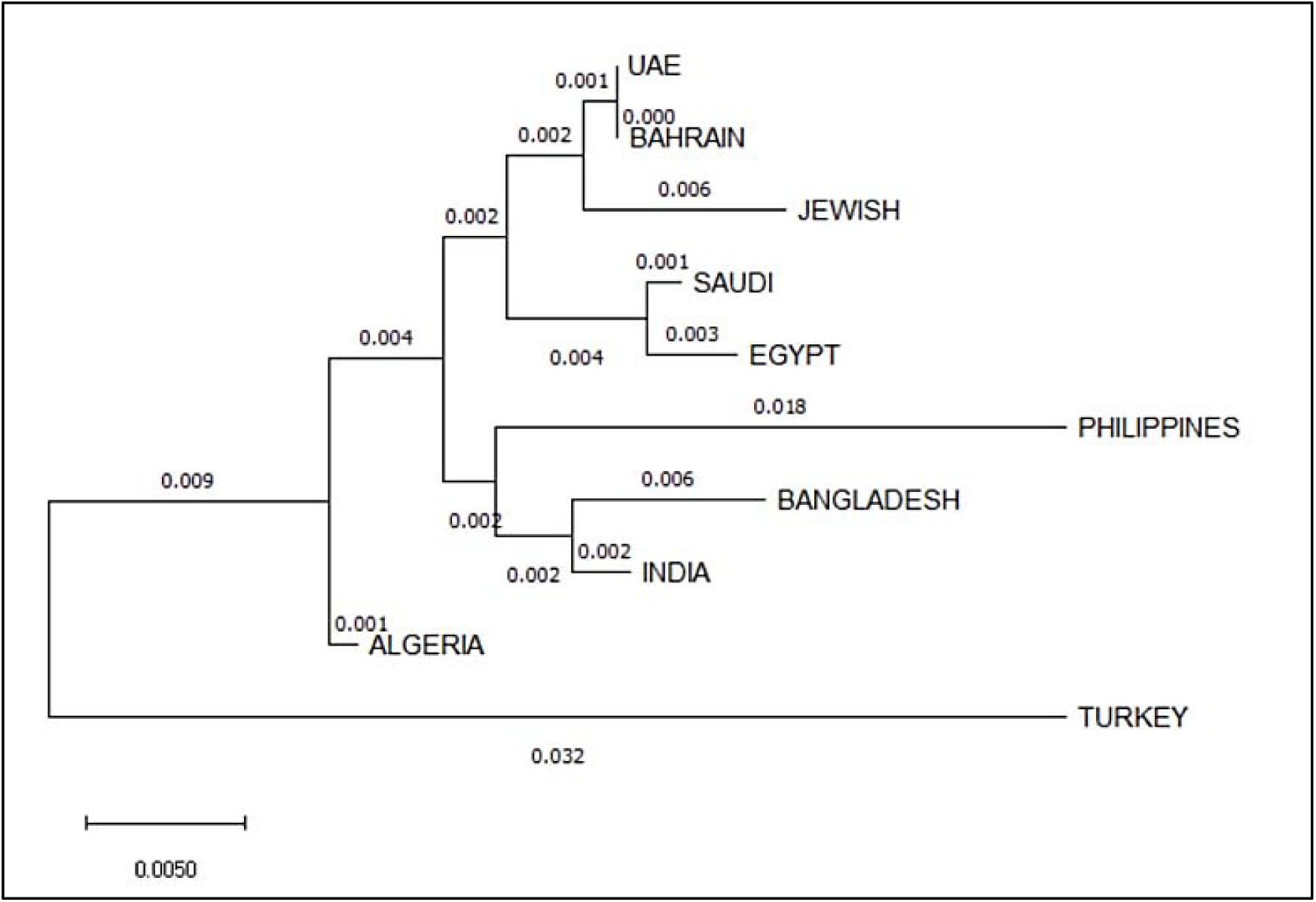
Phylogentic tree using based on Nei’s DA Distances for the 12 X-STR loci estimated among the 13 populations. The nods are bootstrap values derived from 1000 replications. 9 populations including: Saudi (19), Filipino (20), Emiratis (18), Bengali (21), Egyptian (22), Turkish (23), Indian (24), Algerian (25) and Jewish (26).

We have used 9 populations including: Saudi **(19)**, Filipino **(20)**, Emiratis **(18)**, Bengali **(21)**, Egyptian **(22)**, Turkish **(23)**, Indian **(24)**, Algerian **(25)** and Jewish **(26)**. Fst and p-values for allele frequency distribution between Bahraini population and the published groups are shown in **(Table 4)**.

**Table 4.** Nei genetic distance matrix between Bahraini population and other populations. Above the diagonal are p values while below the diagonal are Fst values

|  | Saudi | Emirati | Egyptian | Turkish | Algerian | Jewish | Filipino | Bengali | Indian | Bahraini |
| --- | --- | --- | --- | --- | --- | --- | --- | --- | --- | --- |
| Saudi | 0 | 0.001 | 0.003 | 0.032 | 0.001 | 0.007 | 0.018 | 0.006 | 0.002 | 0.001 |
| Emirati | 0.009 | 0 | 0.004 | 0.002 | 0.002 | 0.002 | 0.002 | 0.002 | 0.002 | 0.003 |
| Egyptian | 0.004 | 0.011 | 0 | 0.009 | 0.003 | 0.004 | 0.002 | 0.004 | 0.004 | 0.012 |
| Turkish | 0.052 | 0.048 | 0.055 | 0 | 0.009 | 0.002 | 0.002 | 0.003 | 0.003 | 0.006 |
| Algerian | 0.013 | 0.01 | 0.015 | 0.042 | 0 | 0.003 | 0.018 | 0.002 | 0.002 | 0.006 |
| Jewish | 0.012 | 0.008 | 0.017 | 0.059 | 0.013 | 0 | 0.002 | 0.002 | 0.002 | 0.002 |
| Filipino | 0.023 | 0.025 | 0.027 | 0.066 | 0.027 | 0.03 | 0 | 0.002 | 0.002 | 0 |
| Bengali | 0.022 | 0.013 | 0.02 | 0.051 | 0.008 | 0.024 | 0.029 | 0 | 0.002 | 0.041 |
| Indian | 0.017 | 0.009 | 0.014 | 0.05 | 0.013 | 0.019 | 0.02 | 0.008 | 0 | 0.018 |
| Bahraini | 0.009 | 0 | 0.012 | 0.051 | 0.012 | 0.007 | 0.027 | 0.016 | 0.012 | 0 |

It is shown that Bahraini and Emirati populations shared the most genetic relatedness than other populations, as the Saudi population considers being the most geographically close population however gave more genetic relatedness with Egyptian population than with Bahraini population. The rest of populations stood further of genetic relatedness with the Bahraini population.

We have also constructed the MDS plot using IBM SPSS Statistics v21.0 Software, and it gave correlating results with the phylogenetic tree. As Bahraini, Emirati and Algerian populations gave the same clusters and Saudi, Egyptian and Jewish in another cluster **(Fig 3)**.

**Figure 3.**
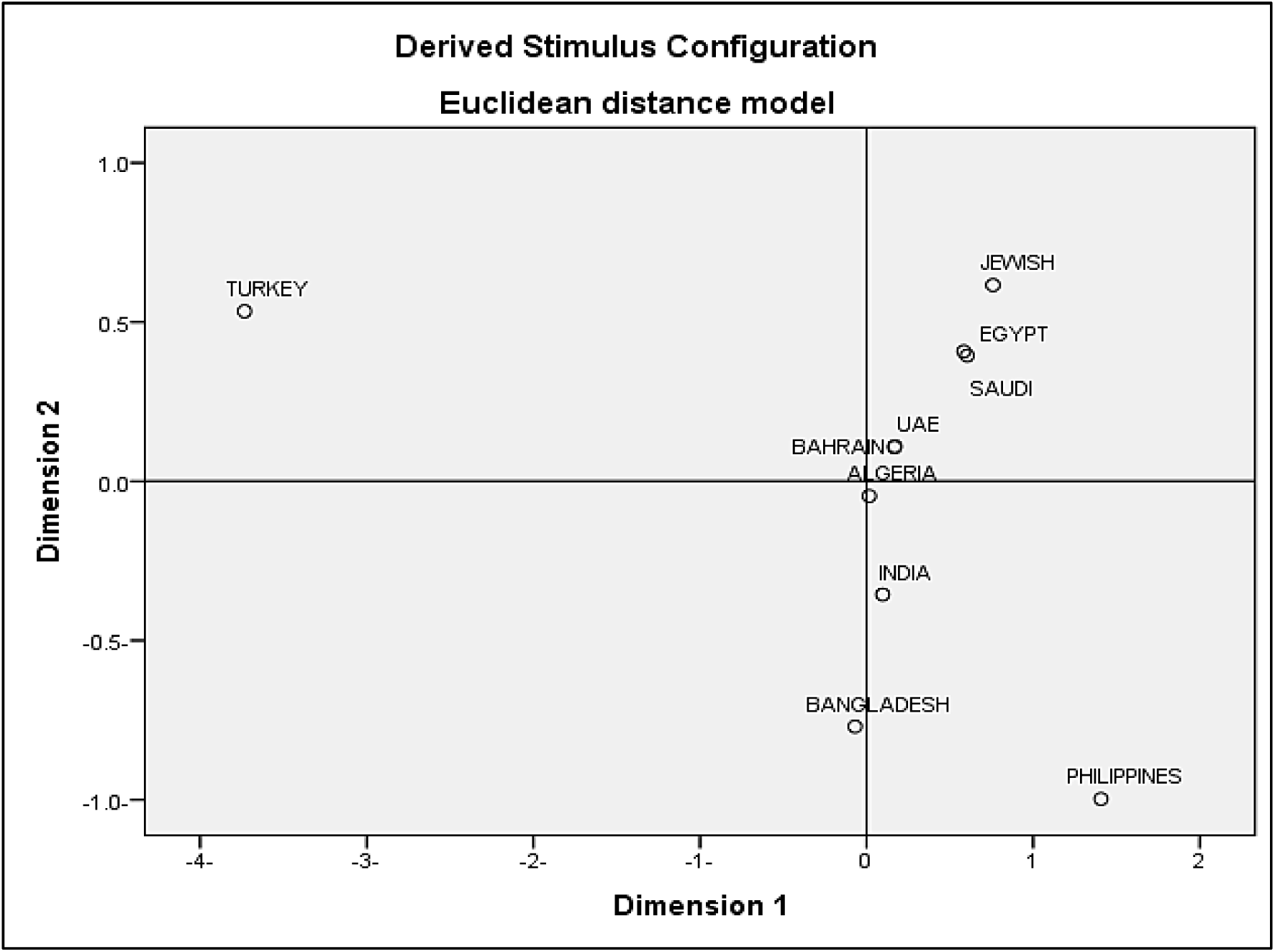
MDS plot constructed between Bahraini population and other populations Stress = .07286 RSQ = .98874

## Discussion

The observed deviation from LD (neglecting the Bonferroni’s correction) could be a result of the high diversity of the Bahraini population or caused by high polymorphism at the same loci investigated loci. This observation are likely to reflect the high level of inbreeding with consanguinity rates in Bahrain, with intra-familial unions accounting for 20–50% of all marriages compared to other Arab countries **(27)**. The PD in correlation with PM supports the high degree of polymorphism between Bahraini individuals.

As shown the significance of LD provided of loci in different LGs, suggesting of using individual allele frequency as well as haplotype allele frequencies for population database and likelihood studies.

We have compared Bahraini population data with other populations according to the available data using the accessible loci **(Table S2)**. It is shown that the Bahraini population shares similar results with the study conducted of Emirati populations using the XSTRs loci. Allele 14 in locus DXS7423 scored the highest frequency for Bahraini, Emirati, Egyptian and Indians populations **(18, 22, 24)** whereas the highest frequency for Filipino and Algerian populations is allele 15 in same locus **(20, 25)**. Turkish population gave the highest allele frequency in allele 13 in HPRTB **(23)**. Jewish and the Saudi populations shared highest frequency with allele 19 in DXS10103 **(19, 26)**. Bengali population showed the highest allele frequency in allele 11 in locus DXS8378 **(21)**.

Regarding the Interpopulation diversity, the phylogenetic tree was constructed based upon the data from the nine populations which were consistent with other population data from the region based upon the FST values obtained. The obtained FST value of Bahrain is <0.0000 which is less than the recommended value for casework statistics of FST < 0.01 **(28)**.

As shown, Bahraini and Emirati populations were more genetically related in terms of phylogenetic tree and MDS plot in contrary of Saudi population which was shown in previous papers published being more genetically related to Bahrain **(4, 5)** This can be explained by studying the origins of the mothers of participants as it wildly affects the X-STRs results.

Once more studies of Arab populations in the region become accessible, it may be more probable to develop a greater understanding of the genetic associations between the different populations for the Arabian Peninsula. Further linkage studies must be conducted to determine if the loci are physically linked.

## Conclusions

In conclusion, this is first study to report the allele frequencies and forensic statistical parameters of the X chromosomal STRs included in the Investigator Argus X-12 QS Kit in a sample population of Bahrain. X-STRs panels can be used for some cases for forensics investigations such as human identification and paternity testing.

It is shown that X-STRs included in the Investigator Argus X-12 kit can be utilized for forensic practice in Bahraini population. Our results demonstrate the importance of analyzing diverse populations using X-STRs markers.

## Acknowledgments

We would like to thank the authorities in General Directorate of Criminal Investigation and forensic Science in Bahrain, namely Mr. Abdulaziz Mayoof Alrumaihi, Mr. Raed Ali Almaeeli and Mr. Mohammed Abdulla Ghayyath for allowing us to utilize the Bahrain forensic Science Laboratory. Also, many thanks to Sabah Nazir and Meshael Ahmed Alqerainees for their technical support. This research did not receive any specific grant from funding agencies in the public, commercial, or not-for-profit sectors.

